# IMMUNOPHENOTYPING AND FUNCTIONAL ANALYSIS OF NK CELL SUBSETS IN *Mycobacterium tuberculosis*-INFECTED INDIVIDUALS

**DOI:** 10.1101/2024.03.16.585339

**Authors:** Carolina S. Silva, Elin Folkesson, Fariba Foroogh, Maia S. Gower, Linn Kleberg, Pengjun Xi, Dhifaf Sahran, Judith Bruchfeld, Margarida Correia-Neves, Gunilla Källenius, Christopher Sundling

**Affiliations:** Division of Infectious Diseases, Department of Medicine Solna, Center for Molecular Medicine, Karolinska Institutet, Stockholm, Sweden; Department of Infectious Diseases, Karolinska University Hospital, Stockholm, Sweden; Department of Laboratory Medicine, Division of Pathology, Karolinska Institutet, Stockholm, Sweden; Life and Health Sciences Research Institute, School of Medicine, University of Minho, Braga, Portugal; ICVS/3B’s, PT Government Associate Laboratory, Braga, Portugal

## Abstract

Infection with *Mycobacterium tuberculosis* remains a global health challenge, with diverse clinical outcomes ranging from latent TB (LTB) infection to active TB disease (ATB). We conducted a comprehensive analysis of NK cell subsets and function in individuals with LTB, ATB, and healthy controls to elucidate their potential association with TB pathogenesis. Peripheral blood mononuclear cells (PBMCs) were isolated from Mtb-infected individuals and analyzed by mass cytometry and flow cytometry. We identified distinct NK cell subsets and evaluated their functional responses to stimulation. Our findings revealed comparable frequencies of total and NK cell subsets across LTB, ATB, and controls. Functional assays demonstrated similar degranulation, cytokine production, and proliferation capabilities among NK cell subsets across the three groups. This study provides insights into the heterogeneity of NK cell responses in TB and highlights the need for standardized methodologies with well-characterized cohorts controlling for donor background. Further investigations are warranted to delineate the specific roles of NK cells in TB immunity and pathogenesis.

## INTRODUCTION

Despite considerable efforts, tuberculosis (TB) is still one of the deadliest infectious diseases worldwide (WHO, 2023). Infection with Mycobacterium tuberculosis (Mtb) is commonly divided into active TB (ATB) or latent TB (LTB), depending on whether the individuals develop clinical symptoms (ATB) or not (LTB) (Pai et al., 2016). However, inhalation of droplets containing Mtb results in different clinical outcomes that essentially depend on the host immune response to the pathogen (Pai et al., 2016). This ranges from elimination of the bacteria, either through the innate or the adaptive immune system, to persistent Infection with long-term control, to disease progression and development of symptoms. Most of the individuals who have had contact with the bacterium retain immunological memory of Mtb antigens but do not progress to an active form of the disease (Behr et al., 2019). This suggests that these individuals can naturally control Mtb growth.

The immune factors underlying the control or progression to TB disease are not clearly defined. T cell-dependent adaptive immunity was thought to be the hallmark of protection in TB, with the production of IFN-γ by CD4^+^ T cells being a key mediator (Cooper, 2009). These cells have been extensively investigated while other immune cell populations have been less explored. However, a protective immune response to Mtb is orchestrated by multiple immune cells and the cytokines secreted by these cells (Nunes-Alves et al., 2014). Natural killer (NK) cells are innate immune cells that target infected host cells via antibody-mediated cytotoxic mechanisms and secretion of proinflammatory cytokines (Mujal et al., 2021). The Mtb bacilli survive in intracellular niches in infected macrophages, potentially indicating an important role of NK cells in controlling the Infection. This is further supported by observations of a unique profile of antibody constant region (Fc) in individuals with LTB associated with enhanced NK cell-mediated Fc effector function (Lu et al., 2016), suggesting a potential interplay between antibodies and NK cells in TB (Li & Javid, 2018).

To better understand the immune cell populations underlying the immune response in various stages of Mtb Infection, we performed comprehensive immune profiling and comparison of NK cells from individuals infected with Mtb who control bacterial growth (LTB) or progress to disease (ATB) and healthy individuals. Moreover, we evaluated the function of NK cells in terms of degranulation, cytokine production, and proliferation.

## METHODS

### Study participants

Participants were recruited between 2018 and 2024. The cohort includes adult (≥18 years) TB patients, contacts to active cases, and IGRA^+^ migrants attending the TB Centre, Department of Infectious Diseases Karolinska University Hospital Stockholm (Supplemental Table 1). Active TB cases were defined upon microbiological (PCR and/or culture) verification; Latent TB Participants were defined as asymptomatic and IGRA^+^ and individuals with recent and remote exposure to Mtb; the IGRA^−^ group includes contacts to active TB cases that were screened as negative in an IGRA test. Participants with ATB, LTB and IGRA^−^ were assessed for standard biochemical measurements

### PBMCs isolation

Venous blood from each participant was collected into EDTA tubes and peripheral blood mononuclear cells (PBMCs) were purified through density gradient centrifugation using Ficoll (Cytiva) and SepMate tubes (StemCell) according to the manufacturer’s instructions. Briefly, whole blood was diluted with an equal volume of PBS supplemented with 2% of heat-inactivated FBS (2% FBS-PBS) and then layered onto 15 mL Ficoll. The cells were centrifuged at 1200g for 10 min with the brake on. The mononuclear cell layer was collected into a new 50 ml tube and resuspended to 45 ml with 2% FBS-PBS. The cells were spun at 300g for 10 min, after which the cells were resuspended into 1-5 mL FBS and counted on a Countess (ThermoFisher Scientific) with trypan blue. The cells were centrifuged at 250g for 10 min and resuspended with freeze media (90% FBS supplemented with 10% DMSO) and placed in a Mr.Frosty freezing container (Sigma) before moving to –80°C overnight followed by long-term storage in liquid nitrogen.

### Analysis of mass cytometry data

Mass cytometry data was repurposed from (Silva et al., 2021). The mass cytometry FCS data files were gated for different cell subsets: CD45^+^ leucocytes, CD45^+^CD3^+^CD20^−^ T cells, CD45^+^CD3^−^CD7^+^ NK cells, CD45^+^CD3^−^HLA-DR^+^ antigen-presenting cells (APCs), using FlowJo™ v10.6.1. The gated populations were exported to new FCS files that were then analyzed using the R-package Cytofkit v1.12.0, which includes an integrated pipeline for mass cytometry analysis (Chen et al., 2016). Cytofkit was run in R-studio version 1.1.463 and R version 3.6.1. For analysis of total leukocytes, 5000 cells were used per sample. For analysis of gated T cells, NK cells, and APCs, 10000 cells were used per sample. Dimensionality was reduced using Barnes-Hut tSNE with a perplexity of 30 with a maximum of 1000 iterations. Clustering was then performed using density-based machine learning with ClusterX (Chen et al., 2016) and cell subsets were identified by visual inspection of marker expression for each cluster.

### Phenotypic and functional assessment of NK cells by flow cytometry

PBMCs were thawed, washed, and resuspended in complete RPMI media supplemented with L-glutamine (2 mM), penicillin (100 U/ml), streptomycin (100 µg/ml), and 10 % heat-inactivated FBS (all from Thermo Fisher Scientific) and then incubated at 37°C over-night. The cells were then plated at 2.5x10^6^ PBMCs/mL and stimulated with either eBioscience™ Cell Stimulation Cocktail (0.081 µM PMA and 1.34 µM ionomycin, Thermo Fisher Scientific), or anti-CD16 (1µg/mL, clone 3G8, Biolegend), IL-12 (10 ng/mL, Biolegend), and IL-18 (100ng/mL, Biolegend) in the presence of brefeldin A and monensin (Thermo Fisher Scientific) for 6 hours. To evaluate cell degranulation, 1µL of anti-CD107a PE-CF594 (BD Biosciences) was added to each well. Stimulated cells were subsequently stained with antibodies targeting surface markers (Supplemental Table 2), according to manufacturers’ instruction. Cells were then fixed and permeabilized using the eBioscience™ FoxP3/Transcription Factor Staining Buffer (Thermo Fisher Scientific) and stained with intracellular antibodies for TNF, IFN-γ, Ki-67 and granzyme B. For the exclusion of dead cells, LIVE/DEAD green fixable viability staining kit (Invitrogen) was used. Cells were acquired using an Aurora spectral flow cytometer (Cytek Biosciences, Fremont, California, USA) and the data was analysed using FlowJo version 10.8.1. NK subsets and TNF, IFN-γ, Ki-67 and granzyme B percentages among NK and NK subsets were evaluated only when the parent population had at least 100 events.

### CMV serology

Cytomegalovirus (CMV) serology was determined using the commercial Human Anti-Cytomegalovirus IgG ELISA Kit (Abcam) following the manufacturer’s instructions.

### Statistical analysis

The histogram and measures of asymmetry and kurtosis were evaluated, and the D’AgosKno and Pearson tests performed to assess the normality assumption for parametric tests; for data not following a normal distribution, non-parametric tests were used. For quantitative data, depending on the underlying distributions, comparisons between groups were performed through independent 2 Brown-Forsythe and Welch ANOVA or Kruskal-Wallis test followed by Dunn’s post-test. Qualitative data was analysed by Chi-squared test. Differences were considered significant when p<0.05. Statistical analyses were performed using Prism9 (GraphPad Software, USA).

### Study approval

Written informed consent was received from all Participants before inclusion in the study, whereby they were pseudo-anonymised. The study was approved by the Regional Ethical Review Board at Karolinska Institutet in Stockholm (approval numbers 2013/1347-31/2 and 2013/2243-31/4) and the Swedish ethical review authority (2019-05438, with addendum 2022-00231-02), in accordance with the Declaration of Helsinki.

## RESULTS

### Investigating the landscape of peripheral blood mononuclear cells in active and latent TB at the single cell level

We performed a comprehensive profiling of the immune cell populations of individuals infected with Mtb, to characterise the immunological status in latent and active TB. We included a total of 30 individuals with ATB, 47 individuals with LTB, and 15 IGRA^−^ controls (Supplemental Table 1). Different numbers of individuals were included in the analyses below and the numbers are indicated in the caption of each figure. To obtain an overview of the blood immune profile at the single-cell level, we first performed a mass cytometry analysis of isolated PBMCs from individuals with ATB (n=5), LTB (n=5) and healthy controls (HC, n=5) (Figure 1). Frozen PBMCs were thawed and left to rest for 24 h, and then a similar number of cells were analysed by mass cytometry (Figure 1A). First, we performed a dimension reduction using t-stochastic neighbour embedding (t-SNE) and clustering analysis of total pre-gated live CD45^+^ cells. From this we identified four main cell types based on expression of immune cell markers: T cells (CD3), B cells (CD20), NK cells (CD3^−^CD7), and myeloid cells (CD33) (Figure 1B-C). T cells were the most abundant cell subset, representing on average 70% of the cells, followed by NK, myeloid cells, and B cells (Figure 1C). The relative frequencies of the identified subsets were similar between the groups (Figure 1D).

**Figure 1.**
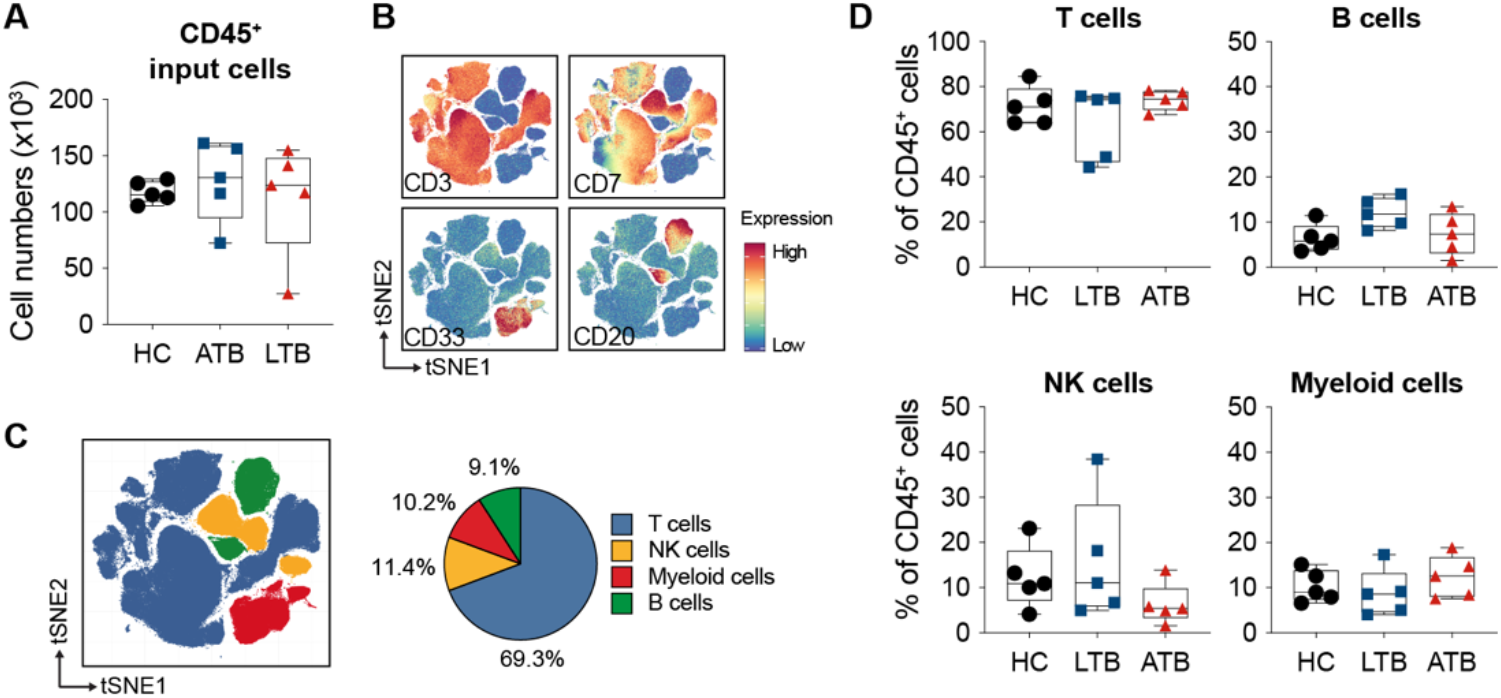
Characterisation of the immune landscape of PBMCs from individuals with ATB and LTB, and HC by mass cytometry. **(A)** Comparison of total CD45+ cells input cells between groups. **(B)** tSNE plots with the expression markers used to characterise the main lymphoid populations. **(C)** tSNE plot of the clustering of CD45+ lymphocytes from all samples together coloured according to cell types (left) and the mean proportions of each cell type (right). **(D)** Comparison of the percentage of T, B, NK, and myeloid cells within total lymphocytes between groups. Statistical differences between groups were analysed using Kruskal-Wallis with Dunn’s post-test (n=5/group).

### Individuals with LTB and ATB have similar percentages of NK cell subsets

To further dissect the immune profile of the leukocytes from the mass cytometry dataset, we analysed the different cell subsets individually. Cells were gated based on CD3, CD7, and HLA-DR expression, and we performed a clustering analysis of each cell subset separately. We identified several subsets among T and APCs, but no differences were found between the subsets (Supplemental Figures 1 and 2). Within NK cells (CD3^−^ CD7^+^), three subsets were identified based on the different expression levels of CD56, CD57, and granzyme B (GrzB) (Figure 2A-B). Although CD56 in combination with CD16 are commonly used as markers to identify NK cell subsets, CD16 is rapidly downregulated in culture, thus not providing a reliable identification. For that reason, CD56 was used in combination with CD57. A subset of CD56^dim^CD57^+^ NK cells, expressing higher levels of GrzB, was found to be increased in individuals with LTB compared to individuals with ATB (Figure 2C).

**Figure 2.**
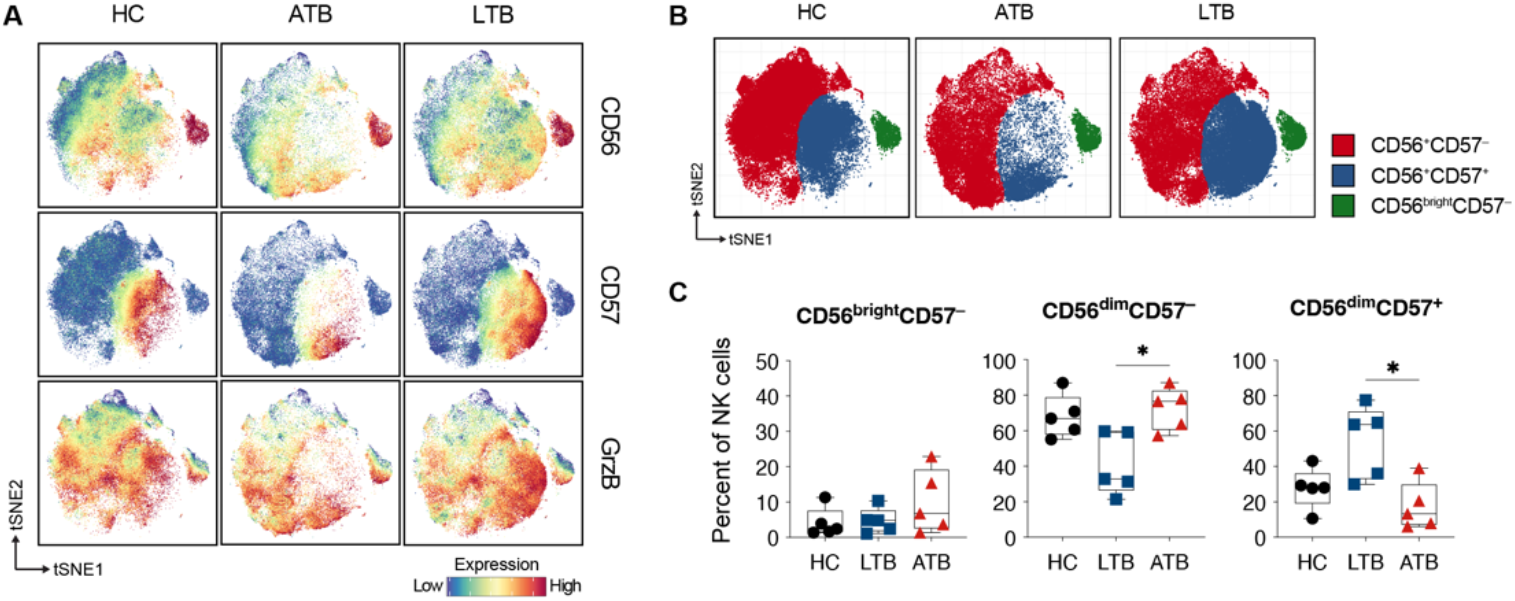
Mass cytometry profiling of NK cells in individuals infected with Mtb shows higher percentages of CD56dimCD57^+^ NK cells in LTB. **(A)** tSNE plots of NK cells with the expression of CD56, CD57, and GrzB in the different groups. **(B)** tSNE plots of the clustering of NK cells across the groups, coloured according to the three main populations identified. **(C)** Percentage of the NK subpopulations within total NK cells from the ATB, LTB, and HC groups. Each dot represents one individual. Statistical differences between groups were analysed using Kruskal-Wallis with Dunn’s post-test (n=5/group). *p<0.05

We then sought to validate the results with a larger number of individuals in each group using flow cytometry. Moreover, since differences in NK cells between endemic and non-endemic countries have been reported (Harris et al., 2020), the control group (IGRA^−^) of the following analyses consists of individuals with similar origin of those from the Mtb-infected groups, and not only Sweden-born individuals (as the HC group). NK cell subsets were determined based on expression levels of CD56 and CD57, similar to the mass cytometry data, and we analysed canonical (CD57^+^NKG2C^−^) and adaptive (CD57^+^NKG2C^+^) NK cell subsets (Figure 3A). Contrary to the observations in the exploratory mass cytometry analysis, no differences in cell frequencies were observed between the groups among the NK cell subset (Figure 3B).

**Figure 3.**
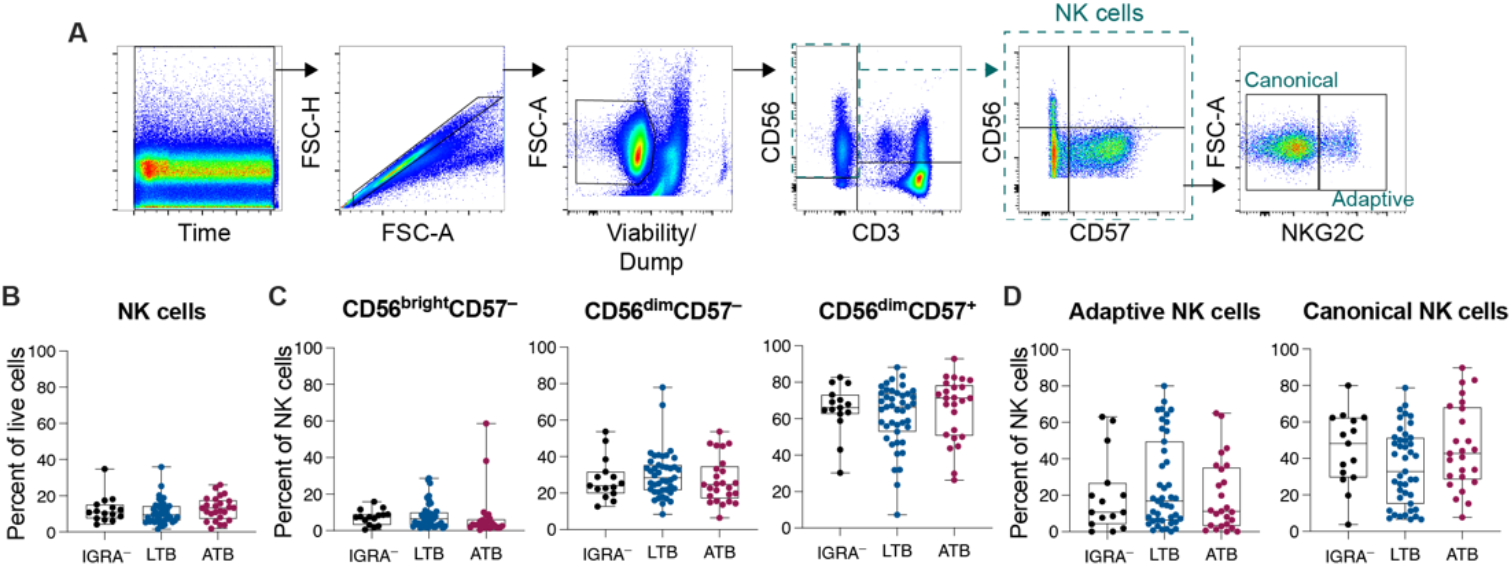
Analysis of NK cells and NK subpopulations by flow cytometry. **(A)** Flow cytometry gating strategy for the evaluation of NK cells’ phenotype. **(B)** Frequency of total NK cells (CD3^−^CD56^+^) out of Live PBMCs. **(C)** Frequency of NK cell subpopulations out of total NK cells based on expression of CD56 and CD57. **(D)** Frequency of canonical (CD57^+^NKG2C^−^) and adaptive (CD57^+^NKG2C^+^) NK cell subsets out of total NK cells. Each dot represents one individual. Statistical differences between groups were analysed using the Brown-Forsythe and Welch ANOVA test. (IGRA^-^=15 LTB=43, ATB=25).

### Individuals with LTB and ATB have similar NK cell function

We next investigated the function of NK cells by evaluating cell degranulation, cytokine production, and proliferation (Figure 4A). Degranulation was assessed by the expression of surface CD107a and intracellular GrzB levels, and proliferation by Ki-67 levels. The cytokines IL-12 and IL-18 are commonly used to stimulate NK cells and to study NK function (Sarhan & Miller, 2019). However, adaptive NK cells (CD56^dim^CD57^+^NKG2C^+^) have low expression of the receptor of these cytokines and responde less to IL-12 and IL-18 stimulation, but are sensitive to stimulation through CD16 (Schlums et al., 2015). Thus, to stimulate all NK cells uniformly, we used IL-12 and IL-18 in combination with anti-CD16. Moreover, to evaluate the overall cell functionality, cells were also stimulated with PMA and ionomycin (PMA/ion).

**Figure 4.**
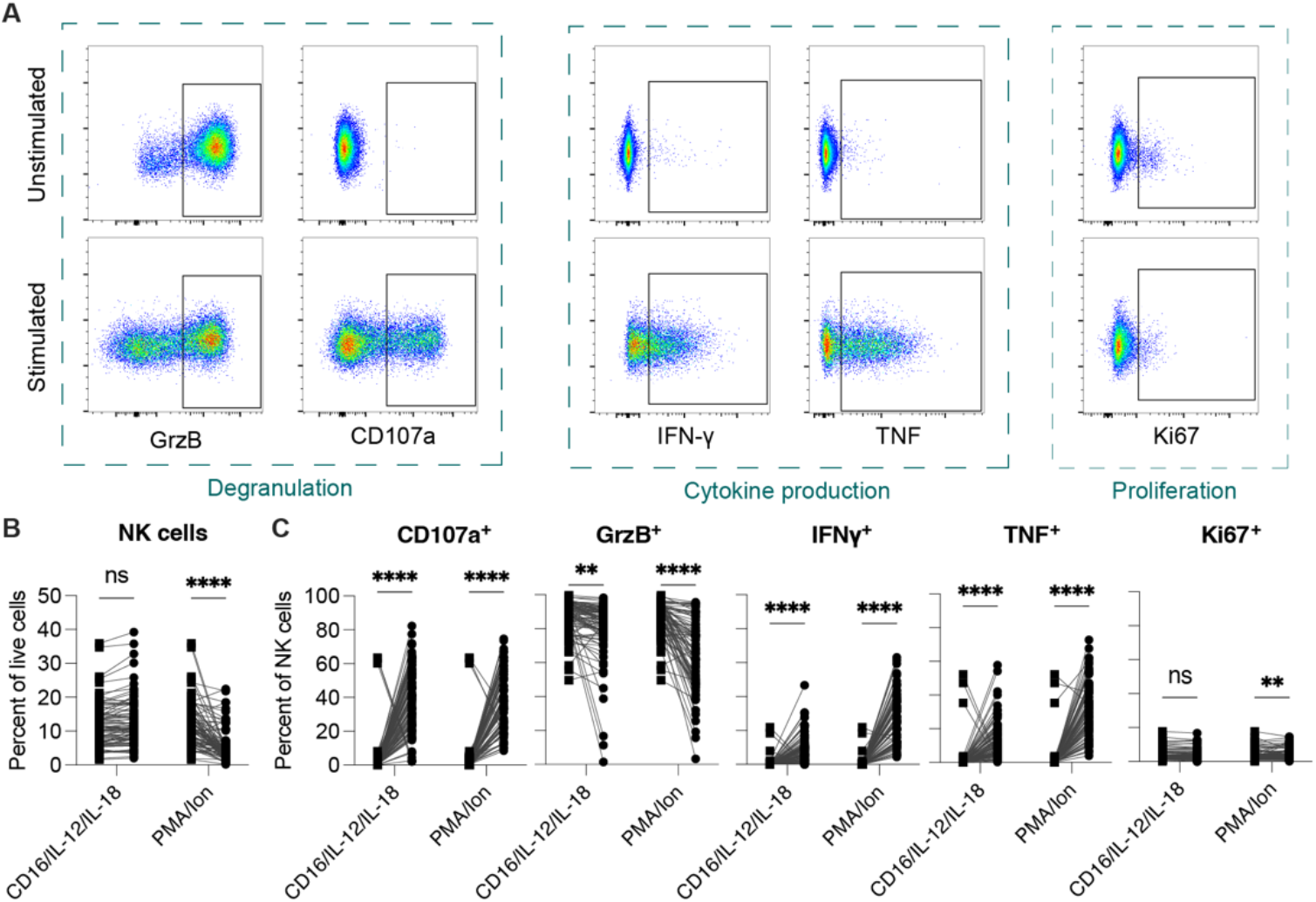
Effect of stimulation on NK cells regardless of the experimental group. PBMCs isolated from individuals with ATB and LTB, and controls were stimulated with α-CD16/IL-18/IL-12 or PMA/ionomycin and cell function was determined by evaluating degranulation (CD107a and GrzB), cytokine production (IFN-γ and TNF) and proliferation (Ki-67). **(A)** Flow cytometry gating strategy for the evaluation of NK cell function. **(B)** Frequency of total NK cells (CD3^−^CD56^+^) out of Live PBMCs. **(C)** Frequency of CD107a, GrzB, IFN-γ and TNF-positive NK cells. Each dot represents one individual. Statistical differences between groups were compared using paired t-test or Wilcoxon matched pairs signed rank test according to data distribution. (n=83/group) **p<0.01, ***p<0.001, ****p<0.0001

Stimulation with PMA/Ion resulted in lower percentages of live NK cells but anti-CD16/IL-12/IL-18 did not affect the frequency of NK cells compared to unstimulated cells (Figure 4B). Both stimulations resulted in degranulation by NK cells as shown by increased expression of CD107a on the cell surface and lower expression of GrzB intracellularly (Figure 4C) (Alter et al., 2004; Prager & Watzl, 2019). Moreover, we observed an increased frequency of cells producing IFN-g and TNF (Figure 4C).

We next analyzed degranulation and cytokine production among the NK cell subsets in response to PMA/Iono (Supplemental figure 3) and anti-CD16/IL-18/IL-12 and in association with Mtb infection status. Overall, there were no differences associated with infection status among total NK cells (Figure 5A) or NK cell subsets (Figure 5B). Since there were no difference between the groups, we next pooled the data and compared NK cell responsiveness between the different NK cell subsets following anti-CD16/IL-18/IL-12 stimulation (Supplemental figure 4). Overall CD56^bright^CD57^−^ cells displayed reduced degranulation, and TNF cytokine production compared with the other CD56^dim^ NK cell subsets. CD56^+^CD57^−^ cells also had slightly reduced degranulation and TNF production. In contrast, Ki67 levels were higher for both the CD56^bright^ and CD56^+^CD57^−^ cells, indicating more recent proliferatiaon. Among the CD56^+^CD57^+^ cells, the adaptive NK cells (NKG2C^+^) had a similar degranulation as the canonical NK cells (NKG2C^−^), but slightly more TNF production. We did not detect any differences in IFN-γ production between the subsets (Supplemental figure 4).

**Figure 5.**
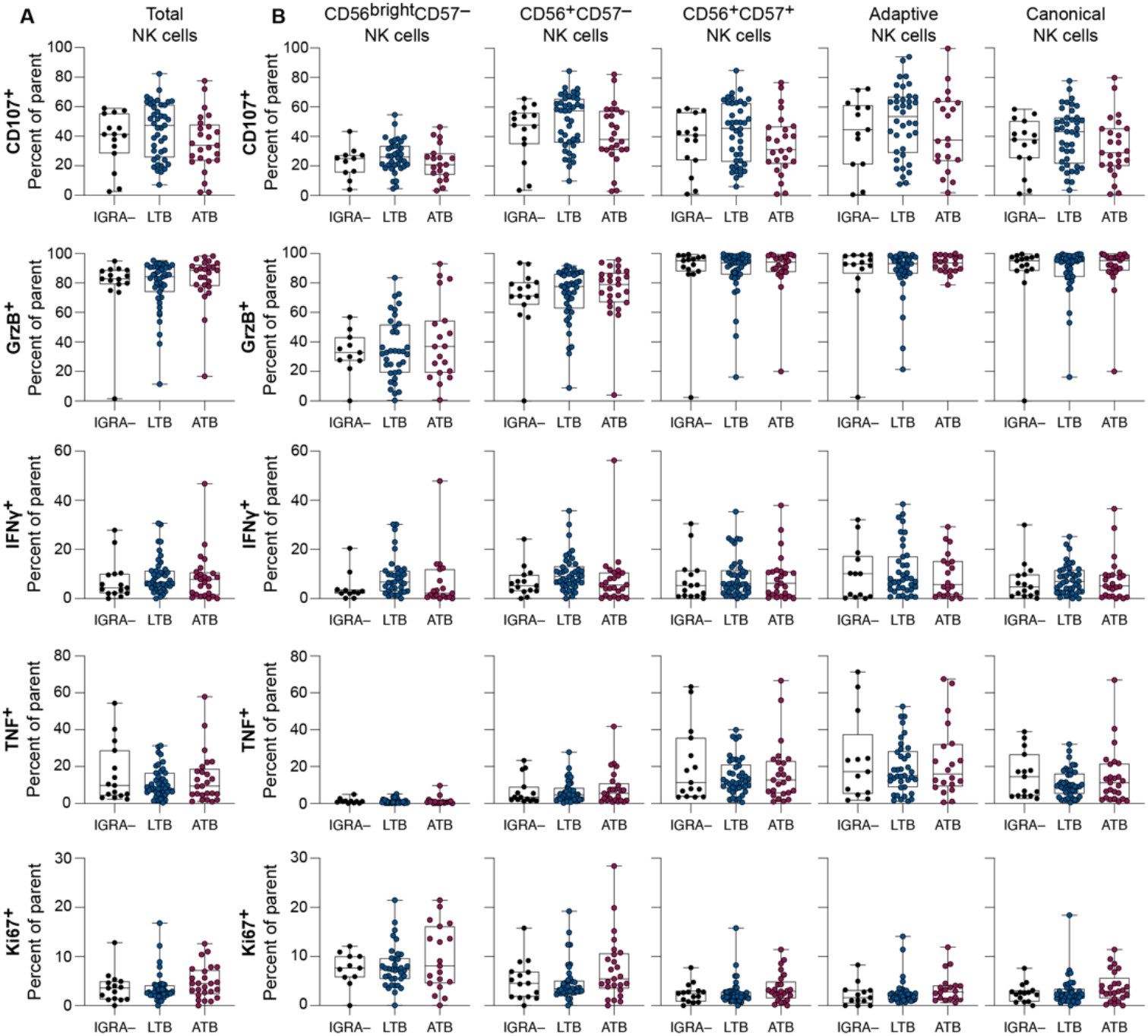
Functional analysis of NK cells and NK subpopulations evaluated by flow cytometry. PBMCs isolated from healthy controls or individuals with ATB, or LTB were stimulated with α-CD16/IL-18/IL-12 and NK cells function was determined by evaluating degranulation (CD107a and GrzB), cytokine production (IFN-γ and TNF) and proliferation (Ki-67). Each dot represents one individual. Statistical differences between groups were analysed using the Brown-Forsythe and Welch ANOVA test. (IGRA-=11-15 LTB=34-43, ATB=19-25).

## DISCUSSION

There is mounting evidence that NK cells play an important role in the immune response to Mtb (Choreño-Parra et al., 2021). In a comprehensive multi-cohort study, Roy Chowdhury et al. observed that the levels of total NK cells decreased during progression from latency to active disease and were restored after anti-TB treatment, suggesting that NK cells could play a role in Mtb infection (Roy Chowdhury et al., 2018). Here we assessed not only the frequency of total NK cells, but also the frequency of the main NK cell subsets in latent and active TB, as well as the function of these subsets in terms of degranulation, cytokine production, and proliferation. We observed that individuals with latent and active TB, as well as IGRA^−^ individuals had similar frequencies of total NK cells. This has also been observed by others (Garand et al., 2018; Harris et al., 2020; Kamolratanakul et al., 2024; Maseko et al., 2023). In contrast, both Roy Chowdhury et al., and Choreño-Parra et al., reported increased total NK cells percentages in individuals with LTB compared to individuals with ATB and healthy controls (Choreño-Parra et al., 2021; Roy Chowdhury et al., 2018), highlighting variation in the NK literature in the TB infection and disease context.

NK cells have different functions, such as cytotoxic and regulatory, and can be divided into different subsets based on the expression of CD56 and CD57 (Fu et al., 2014; Nielsen et al., 2013). In this study, we divided the NK cells into five subsets (CD56^bright^CD57^−^, CD56^dim^CD57^−^, CD56^dim^CD57^+^, adaptive CD56^dim^CD57^+^NKG2C^+^ and canonical CD56^dim^CD57^+^NKG2C^−^) that we then further assessed before and after stimulation. We especially focused on comparing NK cell subset responses between individuals with ATB, LTB, or IGRA^−^ contacts. However, we did not observe any statistical differences associated with infection status or any trends indicating potential differences. This is consistent with a study by Garand et al., where they similarly found no differences in NK subset frequencies between individuals with ATB, tuberculin skin test (TST)^+^ and TST^−^ household contacts (Garand et al., 2018). However, they observed that the frequency of NKG2C^+^ cells within CD16^−^CD56^dim^ NK cells was higher in TST^+^ individuals, compared to TST^−^ and ATB (Garand et al., 2018). We could not evaluate this relatively small subset as we did not include CD16 in our panel and the CD56^dim^CD57^−^NKG2C^+^ cells that we observed would still retain a substantial number of CD16^+^ cells. In a study by Maseko et al., comparing NK subsets between individuals with ATB with and without HIV-co-infection and healthy controls, the authors observed higher frequencies of CD16^+^CD56^neg^ NK cells in ATB/HIV co-infected individuals compared with HIV-negative individuals with ATB and healthy controls but not between ATB and healthy individuals (Maseko et al., 2023). However, in a recent study, it was reported that individuals with LTB who progress to ATB have higher frequencies of CD16^+^CD56^dim^CD57^+^ than those that do not progress (Kamolratanakul et al., 2024).

For many studies on NK cells, there is a relatively large variation between donors in both frequencies/number and responses to stimulation. For some NK cell subsets, we observed a bi-modal distribution, i.e. two separate groups of individuals (Figure 3C-D). This was especially apparent when measuring the frequency of CD56^dim^CD57^+^, adaptive, and canonical NK cells, and it was more pronounced in individuals with LTB and ATB compared to controls. Similar bi-modal distributions have been observations by Garand et al., for CD16^+^ NK cells (Garand et al., 2018) and by Pean et al., for CD56^dim^ NK cells in a study that included TB/HIV-co-infected individuals (Pean et al., 2023). Garand et al., confirmed that age, HIV status, smear grade or chest x-ray grade did not contribute to the bi-modal distribution. We could also confirm that neither age nor sex were associated with the bimodal distribution in our dataset. To further extend the analysis we considered that NK cells are also known to play a large role in the immune response to other infections, such as CMV, where a similar bi-modal distribution was observed in NKG2C^+^ NK cells (van der Ploeg et al., 2023). We therefore measured CMV positivity and antibody levels in our dataset, but neither positivity nor antibody levels could explain the data distribution, leaving this to be further investigated elsewhere.

Studies in non-human primates showed an accumulation of CD27^+^ NK cells in LTB animals, but not in those with ATB (Esaulova et al., 2021). Diedrich et al., evaluated the phenotype and function of NK cells during Mtb-infection in non-human primates and they found no differences in blood and airways NK cell subsets. However, they observed that airway NK cells are more polyfunctional at 4 weeks post Mtb infection, compared to pre-Infection (Diedrich et al., 2023). In non-human primates vaccinated intravenously with BCG, Darrah et al., did not find any significant changes in the frequency of NK cells (Darrah et al., 2020), that has been suggested to correlate with protection (Suliman et al., 2016).

NK cells directly kill infected cells through their cytotoxic activity, but they can also produce cytokines that boost the immune response to intracellular infections, such as Mtb. Roy Chowdhury et al. showed that individuals with LTB had higher levels of circulating cytotoxic NK cells compared with healthy controls (Roy Chowdhury et al., 2018). These changes in NK cells inversely correlated with the lung inflammatory state of individuals with ATB. Garand et al. observed a significant decrease in IFN-γ expression and degranulation in NK cells from TB cases pre-treatment compared to post-treatment (Garand et al., 2018). Here we evaluated NK cell function in LTB, ATB and IGRA^−^ individuals by stimulating total PBMCs with anti-CD16 together with IL-12 and IL-18. We observed that individuals with LTB and ATB, as well as IGRA^−^ had a similar NK cell function in terms of cytokine production and degranulation, which is in contrast with the observations in the above-mentioned studies. However, it is important to note that the experimental setup of the above-mentioned studies differs. Roy Chowdhury et al. evaluated NK cell cytotoxicity by calcein-release assay using target K562 cells while Garand et al. used whole blood instead of isolated PBMCs.

Thus there is a considerable heterogeneity in the scientific literature about the role of NK cells in TB. The different observations of NK cell (subset) frequencies in study groups from the spectrum of Mtb infection and disease might be due to slightly different definitions of experimental groups, cell types and methodology. The diagnosis of latent TB is based on the detection of an adaptive immune response to Mtb antigens. This is done using different tests (either TST or IFN-γ release assays [IGRAs]) and different commercial brands of the same type of test, resulting in discrepancies in the diagnosis of LTB (Young et al., 2005). It is known that only about 5-10% of people diagnosed with LTB will progress to disease in the first years after contact with an index case (Behr et al., 2019). Some of the individuals with LTB may clear the bacteria, suggesting that Mtb control could be very different in Mtb-immunoreactive individuals. In most of the studies mentioned before, there is no information regarding the time since exposure to Mtb. An additional important factor is the control group. Harris et al. showed that TST^−^ individuals in a TB endemic setting differed significantly in NK cell responses when compared with Mtb-naïve healthy controls in the U.S. (Harris et al., 2020). We also observed similar significant differences when comparing our NK cell data with Swedish healthy controls (data not shown), highlighting the importance of controlling for background between the study groups. As indicated above, many studies use slightly different strategies to analyse NK cell subsets, both when it comes to the different protein markers analysed, but also culture conditions and stimulations. This makes it difficult to directly translate results between the studies. We also found, when we tried to assess further NK cell subsets, that the variation between donors made it difficult to obtain enough cells for each condition to draw reliable conclusions. We therefore set a strict requirement of at least 100 events in the parent gate, to reduce the risk of generating biased data. At the same time, this limited the scope of our analysis, potentially making us miss out on relevant differences between the groups. In future studies, it would therefore be valuable to start from more cells, whenever possible. It would also be valuable to standardize NK cell panels that can reliably identify a broad range of NK cell markers representing clinically relevant phenotypes and different functional states, such as indicated here (Cossarizza et al., 2021). Similar approaches would also be welcome for functional assays, as those are also methodologically very variable. In conclusion, our results suggest that NK cell phenotype and function in ATB and LTB are similar. However, considering the large heterogeneity in the literature, it also highlights the importance of further studies using more standardized methodology.

## ACKNOWLEDGEMENTS

The authors acknowledge all the study Participants who generously contributed to the study. We also thank all clinicians, nurses (Anna Löwhagen Welander, Jan Bellbrant, and Monica Modin), and medical students (Jens Hellberg, Mathilda Jakobsson, Lisann Grünewald, and Vera Kjellgren) for helping us with the inclusion of Participants and sample processing. Funding for the study was provided by the Swedish Research Council (2019-0140, 2021-03706, and 2023-01943) to CS and (2019-04663 and 2020-03602) to GK. The Heart-Lung Foundation (20230244) to CS and (20200194 and 20220181) to GK.

## AUTHOR CONTRIBUTIONS

G.K. and C.S. designed the study. C.S.S. and P.X. performed experiments. C.S.S. analysed the data and generated figures and tables. E.F. and J.B. included patients and provided clinical data. C.S.S., F.F., M.G., and L.K. prepared samples. D.S. provided key resources. C.S.S., G.K., and C.S. wrote the first draft of the manuscript. C.S.S., M.C-N., G.K., and C.S. interpreted the results and suggested analysis strategies. All authors contributed to the revision and approved the submitted version.

## SUPPLEMENTARY MATERIAL

**Supplemental Table 1.**
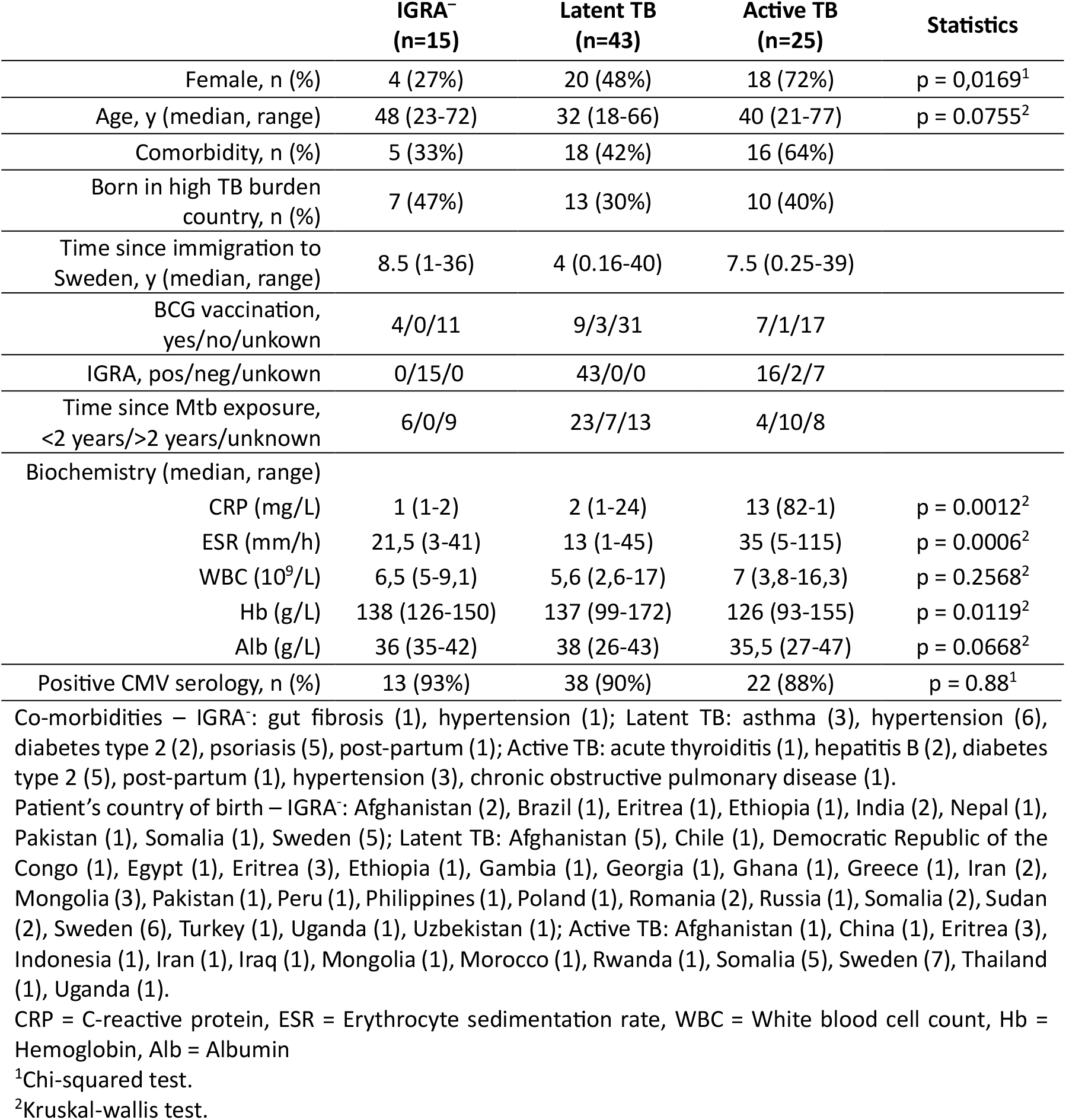
Characteristics of the sample cohort.

|  | **IGRA<sup>-</sup>**<br>**(n=15)** | **Latent TB**<br>**(n=43)** | **Active TB**<br>**(n=25)** | **Statistics** |
| --- | --- | --- | --- | --- |
| Female, n (%) | 4 (27%) | 20 (48%) | 18 (72%) | p = 0,0169<sup>1</sup> |
| Age, y (median, range) | 48 (23-72) | 32 (18-66) | 40 (21-77) | p = 0.0755<sup>2</sup> |
| Comorbidity, n (%) | 5 (33%) | 18 (42%) | 16 (64%) |  |
| Born in high TB burden country, n (%) | 7 (47%) | 13 (30%) | 10 (40%) |  |
| Time since immigration to Sweden, y (median, range) | 8.5 (1-36) | 4 (0.16-40) | 7.5 (0.25-39) |  |
| BCG vaccination, yes/no/unkown | 4/0/11 | 9/3/31 | 7/1/17 |  |
| IGRA, pos/neg/unkown | 0/15/0 | 43/0/0 | 16/2/7 |  |
| Time since Mtb exposure, <2 years/>2 years/unknown | 6/0/9 | 23/7/13 | 4/10/8 |  |
| Biochemistry (median, range) |  |  |  |  |
| CRP (mg/L) | 1 (1-2) | 2 (1-24) | 13 (82-1) | p = 0.0012<sup>2</sup> |
| ESR (mm/h) | 21,5 (3-41) | 13 (1-45) | 35 (5-115) | p = 0.0006<sup>2</sup> |
| WBC (10<sup>9</sup>/L) | 6,5 (5-9,1) | 5,6 (2,6-17) | 7 (3,8-16,3) | p = 0.2568<sup>2</sup> |
| Hb (g/L) | 138 (126-150) | 137 (99-172) | 126 (93-155) | p = 0.0119<sup>2</sup> |
| Alb (g/L) | 36 (35-42) | 38 (26-43) | 35,5 (27-47) | p = 0.0668<sup>2</sup> |
| Positive CMV serology, n (%) | 13 (93%) | 38 (90%) | 22 (88%) | p = 0.88<sup>1</sup> |

Co-morbidities – IGRA: gut fibrosis (1), hypertension (1); Latent TB: asthma (3), hypertension (6), diabetes type 2 (2), psoriasis (5), post-partum (1); Active TB: acute thyroiditis (1), hepatitis B (2), diabetes type 2 (5), post-partum (1), hypertension (3), chronic obstructive pulmonary disease (1).
Patient's country of birth – IGRA: Afghanistan (2), Brazil (1), Eritrea (1), Ethiopia (1), India (2), Nepal (1), Pakistan (1), Somalia (1), Sweden (5); Latent TB: Afghanistan (5), Chile (1), Democratic Republic of the Congo (1), Egypt (1), Eritrea (3), Ethiopia (1), Gambia (1), Georgia (1), Ghana (1), Greece (1), Iran (2), Mongolia (3), Pakistan (1), Peru (1), Philippines (1), Poland (1), Romania (2), Russia (1), Somalia (2), Sudan (2), Sweden (6), Turkey (1), Uganda (1), Uzbekistan (1); Active TB: Afghanistan (1), China (1), Eritrea (3), Indonesia (1), Iran (1), Iraq (1), Mongolia (1), Morocco (1), Rwanda (1), Somalia (5), Sweden (7), Thailand (1), Uganda (1).
CRP = C-reactive protein, ESR = Erythrocyte sedimentation rate, WBC = White blood cell count, Hb = Hemoglobin, Alb = Albumin
<sup>1</sup>Chi-squared test.
<sup>2</sup>Kruskal-wallis test.

**Supplemental Table 2.** Membrane and intracellular NK cell staining panel for flow cytometry.

| Marker | Clone | Fluorochrome | Company | Dilution |
| --- | --- | --- | --- | --- |
| Surface staining |  |  |  |  |
| CD107a | H4A3 | PE-CF594 | BD Biosciences | 1 μL/well |
| NKG2C | S19005E | PE | BioLegend | 1:80 |
| CD3 | SK7 | APC-Fire810 | BioLegend | 1:40 |
| CD19 | SJ25C1 | BV510 | BD Horizon | 1:20 |
| CD14 | MϕP9 | BV510 | BD Horizon | 1:40 |
| CD56 | NCAM16.2 | BV786 | BD Horizon | 1:160 |
| CD57 | TB01 | eFluor450 | Invitrogen | 1:40 |
| Viability staining |  |  |  |  |
| LIVE/DEAD fixable dye | - | Aqua | Invitrogen | 1:600 |
| Intracellular staining |  |  |  |  |
| Ki-67 | SolA15 | AlexaFluor532 | Invitrogen | 1:40 |
| GrzB | QA16A02 | APC | BioLegend | 1:160 |
| IFN-γ | 4S.B3 | BV711 | BD Horizon | 1:40 |
| TNF | Mab11 | BV650 | BD Horizon | 1:40 |

**Supplemental Figure 1.**
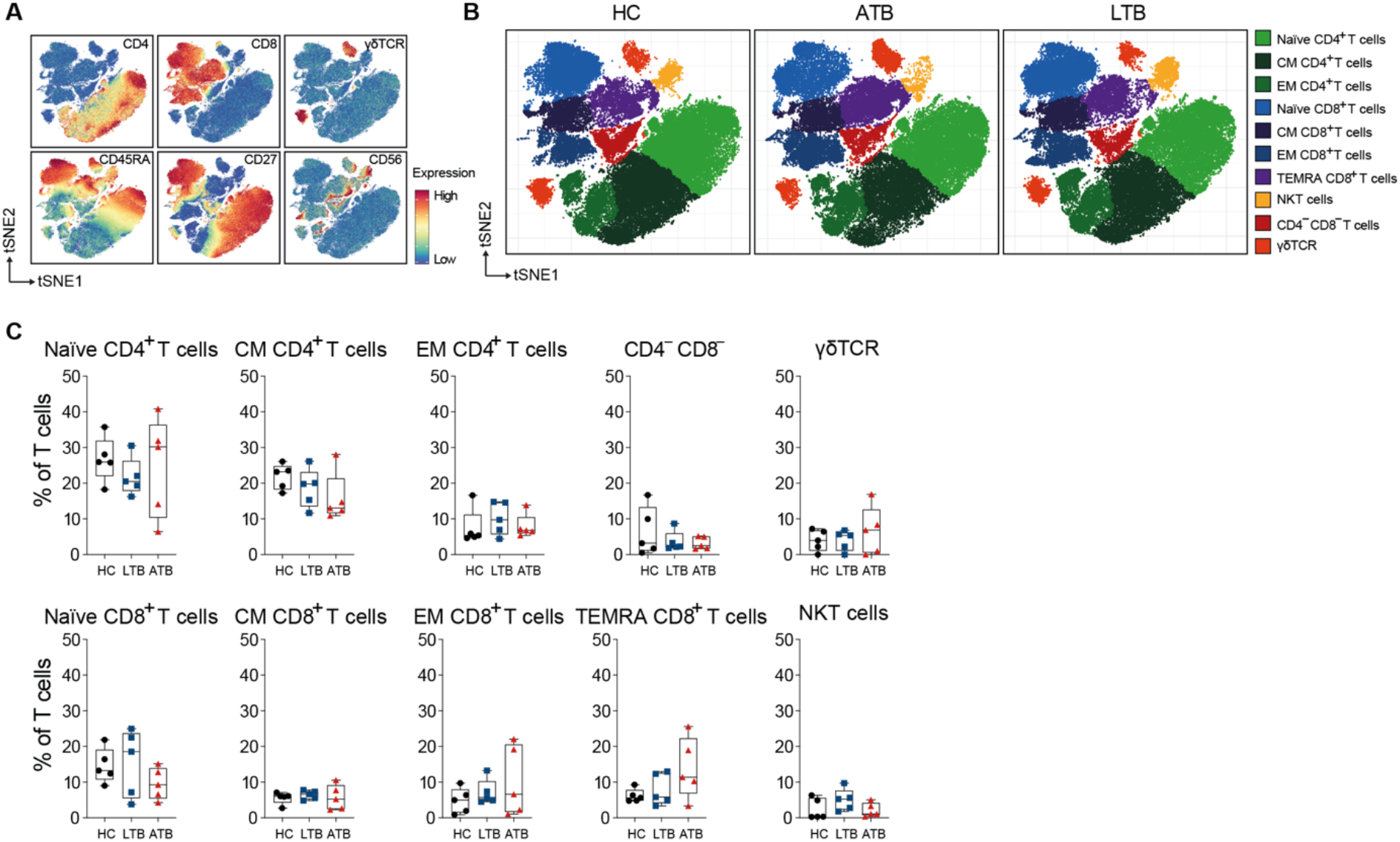
Mass cytometry profiling of T cells in individuals with ATB and LTB, and healthy controls. **(A)** tSNE plots with the expression markers used to characterise the main T cell subsets. **(B)** tSNE plots of the clustering of T cells across the groups, coloured according to the main populations identified. **(C)** Percentage of the T cell subpopulations out of total T cells from the ATB, LTB, and HC groups. Each dot represents one individual. Statistical differences between groups were analysed using Kruskal-Wallis with Dunn’s post-test (n=5/group).

**Supplemental Figure 2.**
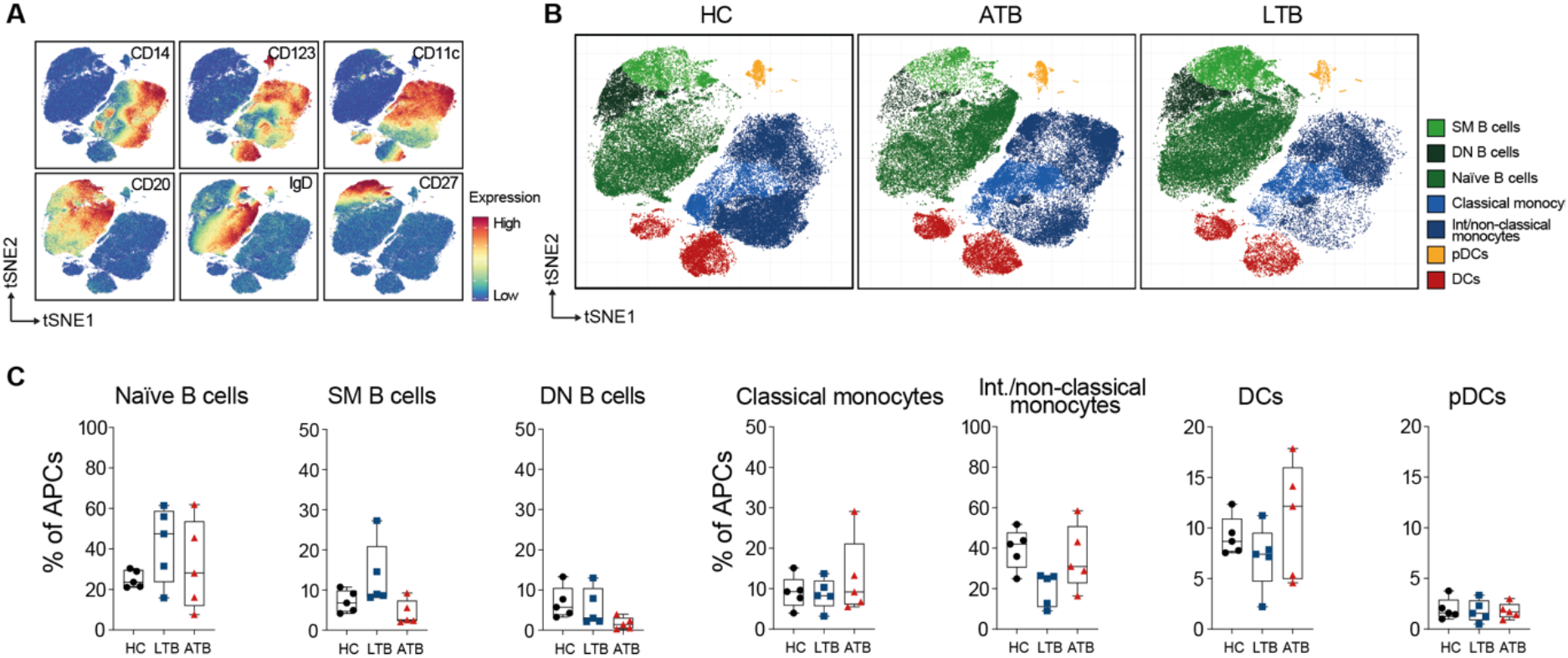
Mass cytometry profiling of APCs in individuals with ATB and LTB, and healthy controls. **(A)** tSNE plots with the expression markers used to characterise the main APC subpopulations. **(B)** tSNE plots of the clustering of APCs across the groups, coloured according to the main populations identified. **(C)** Percentage of B cell and monocyte subsets, DCs, and pDCs out of total APCs from the ATB, LTB, and HC groups. Each dot represents one individual. Statistical differences between groups were analysed using Kruskal-Wallis with Dunn’s post-test (n=5/group).

**Supplemental Figure 3.**
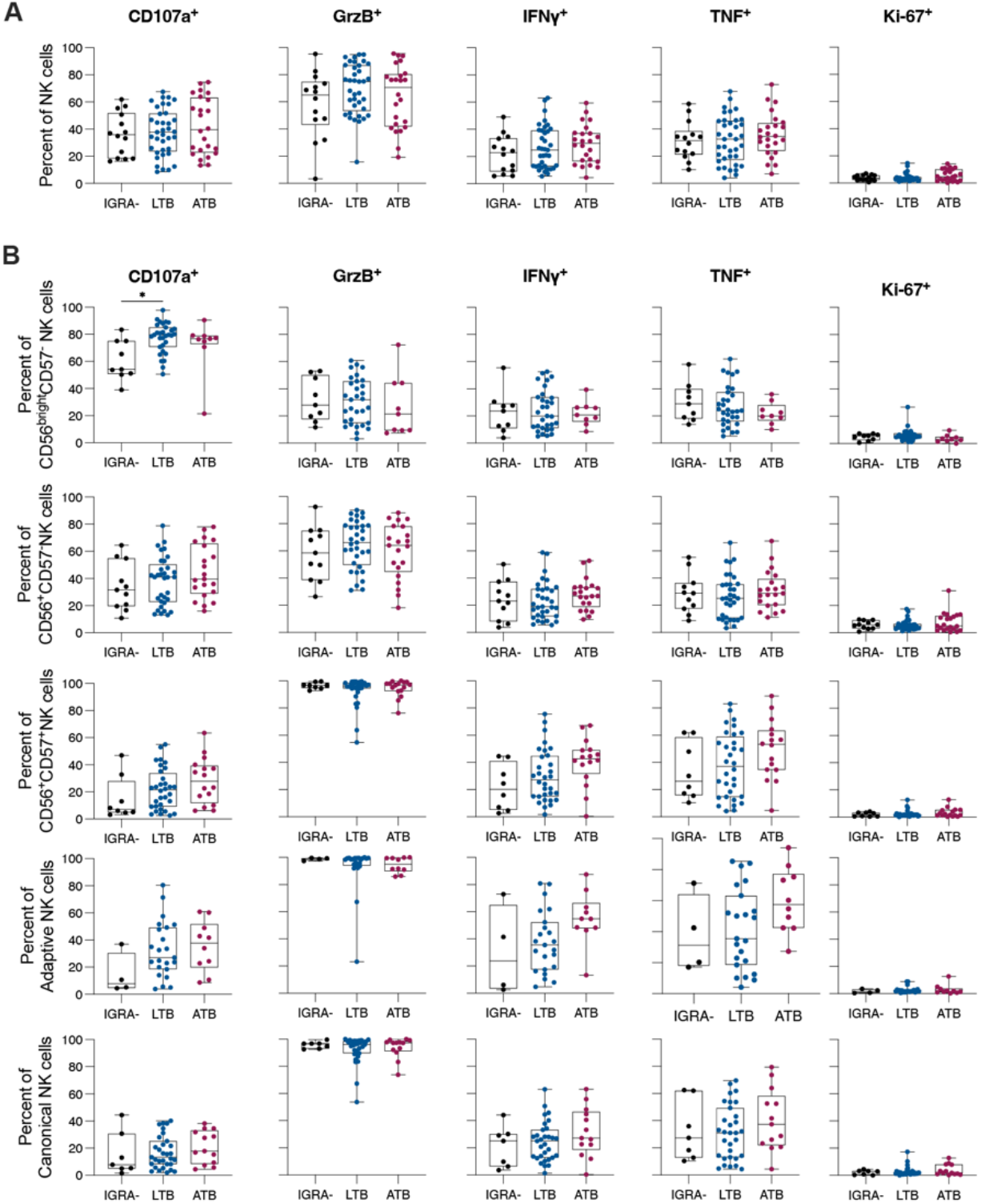
Functional analysis of NK cells and NK subpopulations evaluated by flow cytometry. PBMCs isolated from healthy controls or individuals with ATB, or LTB were stimulated with PMA/Ion and NK cells function was determined by evaluating degranulation (CD107a and GrzB), cytokine production (IFN-γ and TNF) and proliferation (Ki-67). Each dot represents one individual. Statistical differences between groups were analysed using the Brown-Forsythe and Welch ANOVA test. (IGRA-=4-15 LTB=31-43, ATB=9-25).

**Supplemental Figure 4.**
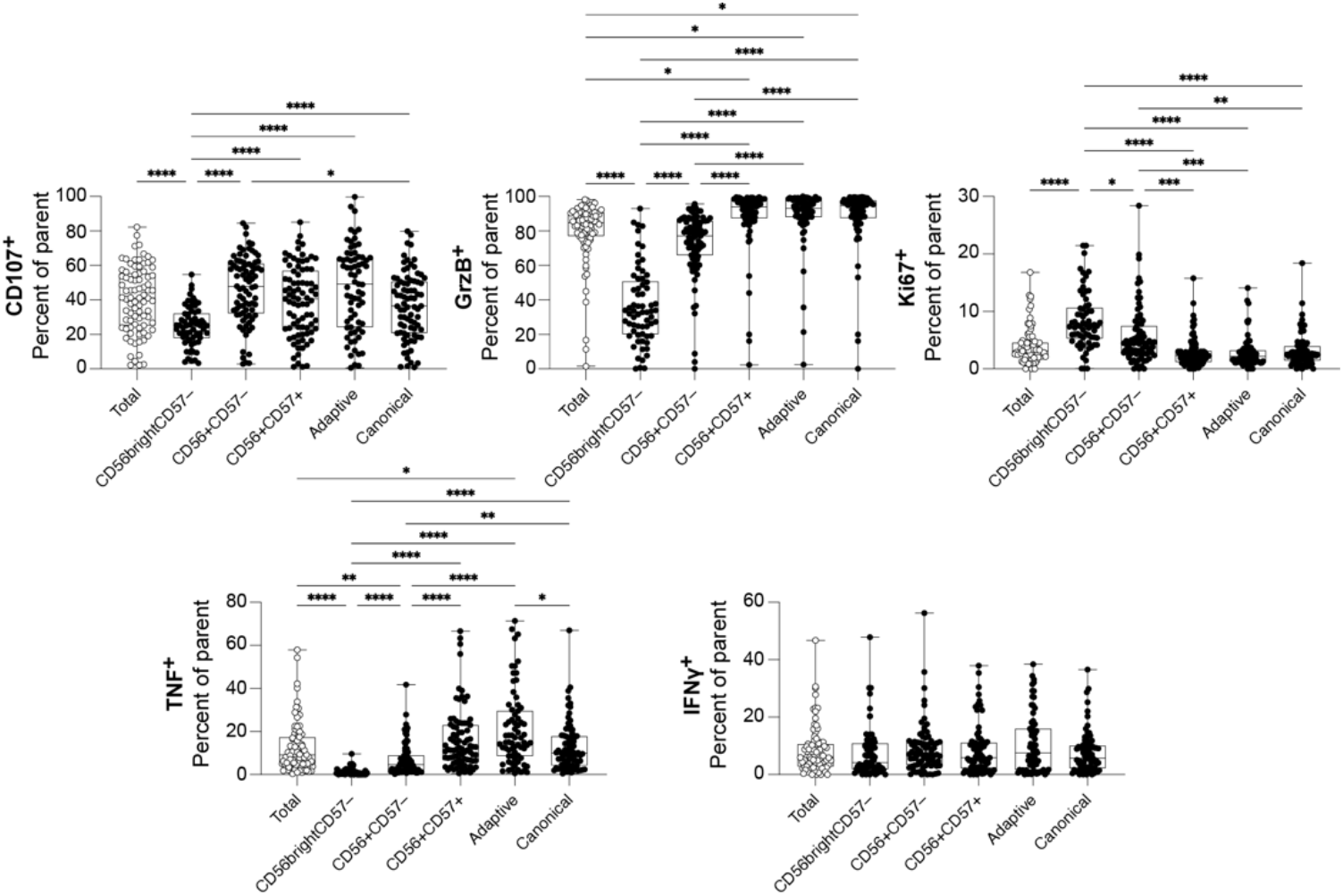
Comparison of NK cells and NK subpopulations followed stimulation with α-CD16/IL-18/IL-12 regardless of the experimental group. PBMCs isolated from healthy controls or individuals with ATB, or LTB were stimulated with α-CD16/IL-18/IL-12 and NK cells function was determined by evaluating degranulation (CD107a and GrzB), cytokine production (IFN-γ and TNF) and proliferation (Ki-67). Each dot represents one individual. Statistical differences between groups were analysed using the Brown-Forsythe and Welch ANOVA test. (n=66-83). *p<0.05, **p<0.01, ***p<0.001, ****p<0.0001

